# Little heterosis found in Diploid Hybrid Potato

**DOI:** 10.1101/2021.11.10.468131

**Authors:** James R. Adams, Michiel E. de Vries, Chaozhi Zheng, Fred A. van Eeuwijk

## Abstract

Hybrid potato breeding has become a novel alternative to con-ventional potato breeding allowing breeders to overcome intractable barriers (e.g. tetrasomic inheritance, masked deleterious alleles, obligate clonal propagation) with the benefit of seed-based propagule, flexible population design, and the potential of hybrid vigour. Until now, however, no formal inquiry has adequately examined the relevant genetic components for complex traits in hybrid potato populations. In this present study, we use a two-step modelling approach to estimate the relevant variance components to assess the magnitude of the general and specific combining abilities (GCA and SCA, respectively) in diploid hybrid potato (DHP). SCA effects were identified for all yield components studied here warranting evidence of non-additive genetic effects in hybrid potato yield. However, the estimated GCA effects were on average two times larger than their respective SCA quantile across all yield phenotypes. Tuber number GCA’s and SCA’s were found to be highly correlated with total yield’s genetic components. Tuber volume appearing under-selected in this population. The prominence of additive effects found for all traits presents evidence that breeders can perform hybrid potato evaluation using the mid-parent value alone. Heterotic vigour stands be useful in bolstering simpler traits but this will be very dependent on the target market of a population. This study represents the first diallel analysis of its kind in diploid potato using material derived from a commercial hybrid breeding programme.

## 1 Introduction

Potato (*Solanum tuberosum*), a plant species once isolated to the continents of southern and central America, is now a crop that spans over 17 million hectares of crop-land worldwide (FAOSTAT 2021). It is the most prominent of non-cereals and is considered by many a major keystone in guaranteeing food security for both local and global communities. Prized for their edible starch-rich tubers, potato meets the demand of several key industries including the fresh, processing, and seed potato markets with a global gross value of 140.5 billion USD as of 2019. As a field crop, potato has a competitive harvest index of 0.85 (in contrast to 0.4-0.6 seen in other crops) in conjunction with a high water productivity (Hay 1995; Lutaladio and Castaldi 2009). There is also ample variation in potato’s tuberization timing requiring as few as 75 days from planting to harvest. All the above make potato a highly productive crop amenable to a variety of cropping systems capable of supplying valuable starch with less agronomic input.

Despite potato’s growing economic and societal importance, rates of crop improvement in complex traits have not kept in step with other major crops over the past century (Douches et al. 1996; Hirsch et al. 2013). Reasons for these deficits in genetic gain are numerous (e.g. market segmentation, large inventory of quality traits, etc.) but many of them stem from the complexities of potato’s evolution and domestication. Potato’s tetraploidy is an oft-cited stumbling block for breeders impeding the ability to fix beneficial loci, and conversely, remove deleterious sites harboured across the genome (Lian et al. 2019; Zhang et al. 2019). Not only does polyploidy mask deleterious loci from traditional forces of selection, but it also impacts the length of time for site fixation even under genetic drift leading to greater maintenance of heterozygosity over time (Bartlett and Haldane 1934). Taken together with a very strong self-incompatibly mechanism, potato could best be described as a fortified heterozygous out-crosser. These biological realities shaped potato breeding from the beginning with breeders conducting crosses between promising heterogeneous individuals followed by the evaluation of large nurseries in search for decent complementation (Simmonds 1979). These F1 nurseries were then subjected to as many as eight subsequent rounds of clonal selection until only elite candidates were left (John E. Bradshaw 2017). While in some ways this method of clonal breeding is quite efficient (all genetic factors are effectively *fixed* at the creation of the F1), it is widely known for being a long process from generation of the nursery to variety release. Because the success of clonal breeding is highly dependent on the generation of enough novel genotypes in the F1, it takes as many as 9 years to sufficiently bulk tubers in conjunction with applying appropriate selection pressure (Bryan 1981; G. C. Tai and Young 1984). It should be noted that while there have been proposals to optimise conventional clonal breeding (Neele, Nab, and Louwes 1991; J. E. Bradshaw, Dale, and Mackay 2003), many of the aforementioned issues are simply implicit to breeding tetraploid potato.

One solution to this comprehensive set of challenges is the adaptation of potato from a tetraploid clonal crop to that of a diploid inbred-hybrid one, an idea which has existed in some form for over 60 years (Hougas and Peloquin 1958). The benefits of such a change, if possible, are manifold; Diploids only take one generation to half their heterozygosity in contrast to an autotetraploid which takes upwards of four generations making the production of pure-breeding lines plausible in the former. As an extension of this, superior genetic performance in the final marketed variety is not dependent on a single crossing event that generated the original F1 (as it is in conventional clonal breeding) but is accomplished through multiple stages (e.g. parental pool improvement, parental line development, hybrid crossing, etc.). This is not to mention other logistical niceties such as the ease of producing & storing true potato seed over vegetative propagule (Cock 1983; Pallais 1991; Thomas-Sharma et al. 2016). Despite the potential of diploid potato, however, it was not until the cusp of the 21st century that it became broadly feasible. Many picked up on the work of (Hosaka and Hanneman 1998) and began the process of generating self compatible populations through the use of *Sli*. (Lindhout et al. 2011) was one of the first to confirm the commercial viability generating diploid potato populations capable of inbreeding using a *Sli* donor. Several subsequent studies not only corroborated that inbred populations in diploid potato were possible (Alsahlany et al. 2021), but hybrids generated from these populations resulted in a crop that could compete in the same space as tetraploid potato (J. Stockem et al. 2020; Zhang et al. 2021). While diploid hybrid potato (DHP) populations are now extant across the world, there is at this time little known about the genetic components controlling complex traits as DHP is still a young hybrid crop. Understanding this is an imperative for potato breeders in order to structure breeding programmes which are able to best exploit the genetic variation available to DHP.

We set out to inspect tuber yield in a large DHP test-cross population. To do this, we evaluated total yield along with two of its simpler yield components, average tuber volume and total tuber number, and partitioned their underlying genetic effects into additive and non-additive components. This trait and genetic decomposition was done to inspect two broad questions: (1) Which tuber phenotypes in this population were responsible for the variation seen in total yield and (2) are these yield phenotypes primarily under the control of additive or non-additive gene action? This latter question holds particular weight as it gives insight on where the focal point of DHP breeding should lie. We put forward a two-part modelling approach to utilise intra-block information to estimate the general and specific combining abilities of our hybrid parents and crosses, respectively. Our study presents the first diallel study in DHP using highly-inbred parents derived from a commercial breeding programme.

## 2 Materials & Methods

### 2.1 Crosses and Trials

A panel of 400 inbred parents were selected and crossed according to distinct selection criteria related to fertility and agronomic traits yielding 806 successful F1 crosses. In the Spring of 2019, all hybrid TPS were sown in trays and grown out in a greenhouse. In May, all seedlings were transplanted at stage 1.0 development (See (Kacheyo et al. 2021)) into two field trials located in the Dutch towns of Est and Heelsum. Both trials utilised a double ridge design with eight plants per ridge with a total of sixteen plants per plot; this design was chosen to minimize within-plot variation while reducing planting costs across each trial (.J E. Stockem et al. 2021). Plots were organised in an augmented randomised complete block design with two blocks and three internal controls used across each block. All 806 F1 hybrids were planted in Heelsum with a subset of 608 hybrids planted in Est. Trial conditions were similar with regards to field management and scoring. One distinguishing factor between trials were their soil conditions with Est being characterised by a light clay composition and Heelsum conversely by distinctively sandy conditions (see Table 1). Both trials were conducted through the summer until haulm killing in early September followed by subsequent harvest two weeks later. All hybrids were scored by several criteria including relevant yield related traits which are our primary focus for this study, i.e., total yield, tuber number and tuber volume. Total tuber number (TN) was measured as the total number of tubers harvested from a given plot of sixteen plants. Tuber volume (TV) was calculated using an average over all tubers harvested per plot using a tuber’s length, width, and depth dimensions to calculate volume using an ellipsoid approximation. Lastly, total yield (TY) was calculated through a transformation of the total tuber weight of a plot to estimate the approximate yield in terms of *Mg* · *Ha*^−1^. These traits were all phenotyped via an automated pipeline described in (J. Stockem et al. 2020).

**Table 1.**
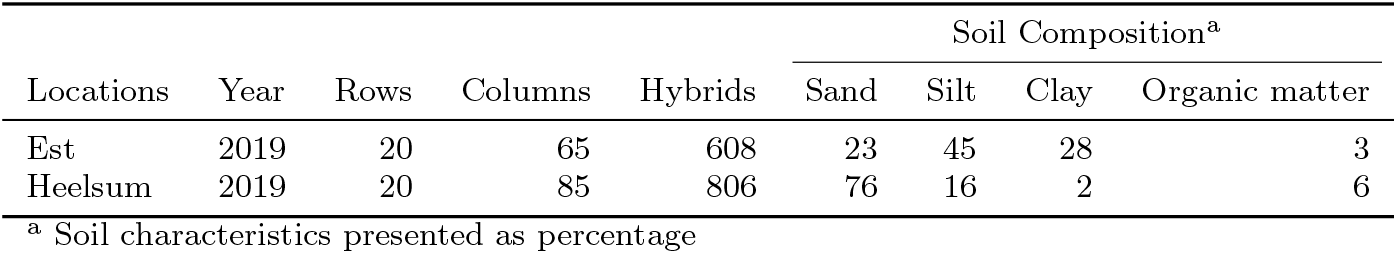
Agronomic properties of the screening trials conducted in Est and Heelsum

| Locations | Year | Rows | Columns | Hybrids | Soil Composition<sup>a</sup> |  |  |  |
| --- | --- | --- | --- | --- | --- | --- | --- | --- |
|  |  |  |  |  | Sand | Silt | Clay | Organic matter |
| Est | 2019 | 20 | 65 | 608 | 23 | 45 | 28 | 3 |
| Heelsum | 2019 | 20 | 85 | 806 | 76 | 16 | 2 | 6 |

<sup>a</sup> Soil characteristics presented as percentage

### 2.2 Spatial Models

This present study used a two-step modelling approach whereby each field trial was modelled separately accounting for factors like field design, control effects, and spatial heterogeneity allowing for extraction of spatial trends and de-trended phenotype data. This was followed by modelling of genetic components for each phenotype. The first step was accomplished by smoothing field effects into local and global trends using two-dimensional penalised splines. This was performed using the Spatial Analysis of Field Trials with Splines (SpATS) library available through the comprehensive R archive network (CRAN) (Rodríguez-Álvarez et al. 2018). While many attested methods capable of handling geo-spatial trends exist, the spatial smoothing approach offered through SpATS was chosen for a few reasons. Often, genetic modelling requires the creation of many spatial models with different spatial structures in order to identify the most satisfactory spatial model. SpATS, conversely, does not follow this procedure and is capable of offering comparable genotype estimates with the best traditional spatial model (Velazco et al. 2017). Along with this, SpATS provides a number of internal methods allowing for intelligible and simple model diagnostics to help elucidate the predominate factors for a given field trial. We chose to model field dimensions using SpATS’ PS-ANOVA method which is capable of taking the bivariate surface and decomposing it into multiple spatial components all defined by one smoothing parameter (Lee, Durbán, and Eilers 2013). The resultant model equation is then:

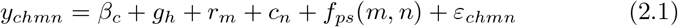

where *β*_*c*_ is a fixed effect for whether the hybrid was a control, *g*_*h*_ is a random effect for the *h*^*th*^ hybrid, *m* and *n* are random effects for row *m* and column *n*. The row and column coordinates were also used within *f*_*ps*_ which represents the two-dimensional penalised-spline function. The PS-ANOVA was parametrised using 19 & 83 internal knots for Heelsum and 19 & 63 internal knots for Est. The large number of internal knots resulted in longer computational time, but were selected to mirror the number of plots along each row and column for each trial. Third degree polynomial B-splines with second degree penalties were used for all spatial models, from which, spatial trends were derived and then subsequently used to de-trend the phenotype data for each trait:

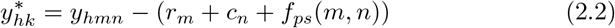

where *y*\* represents the corrected phenotype with systematic spatial trends removed. Each spatial trend was presented as a percentage deviation from the trial mean (see Figures 1 & 2). Along with this, every spatial model was evaluated on the basis of effective and nominal dimension number estimated for each model effect. These were used to evaluate the number of parameters estimated for smoothing and random terms. Taking the ratio between effective and nominal dimensions for random hybrid effects have the benefit of being interpreted as a generalised heritability where the effective dimension number for a hybrid genotype effect (the trace of its hat matrix) is divided by its nominal dimension (the rank of its design matrix) allowing for a direct assessment of genetic variation exhibited within a field trial (Oakey et al. 2006).

**Fig. 1.**
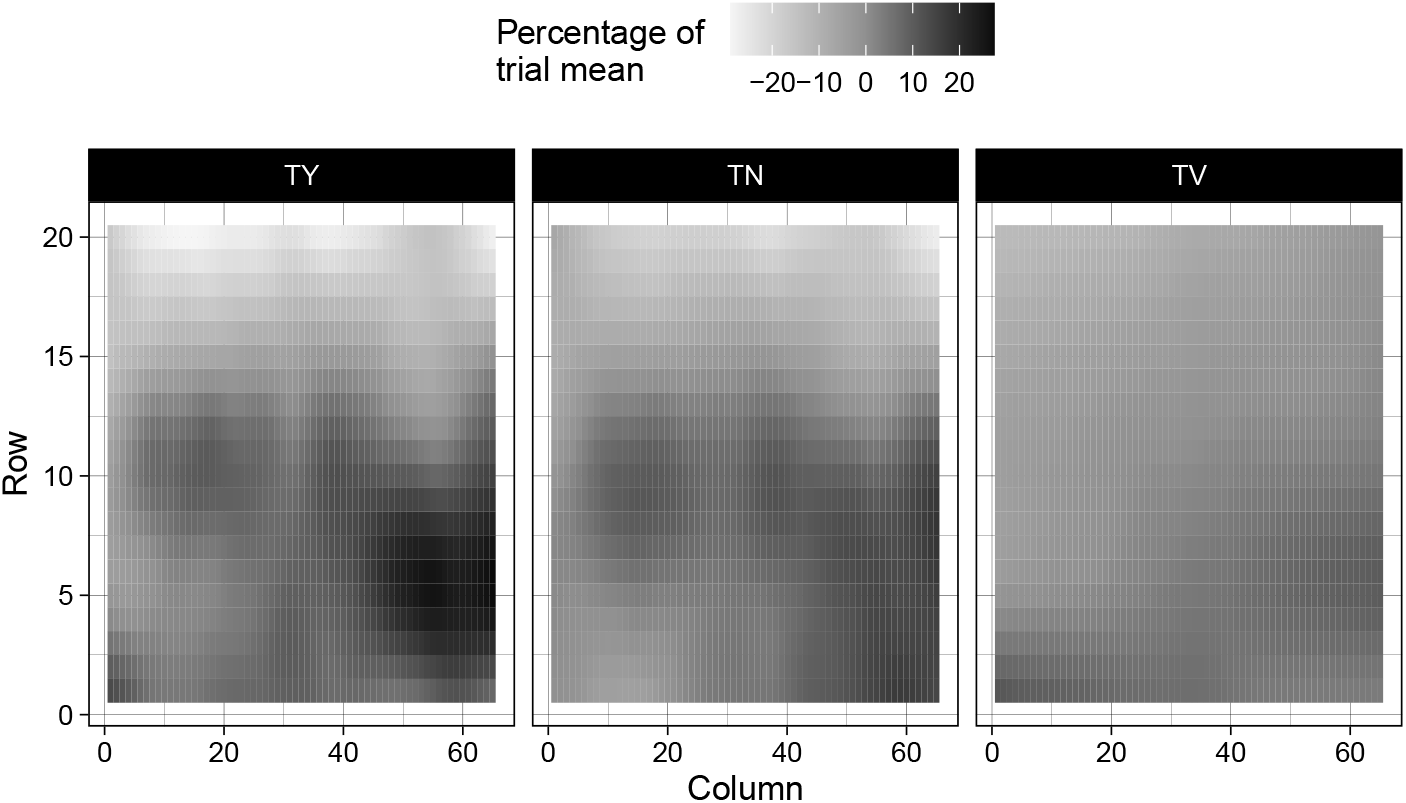
Estimated spatial trends scaled by trial mean for total yield (TY; 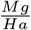) , tuber number (TN; Number of tubers per plot), and tuber volume (TV; Average *cm*^3^ per plot) within the Est screening trial

**Fig. 2.**
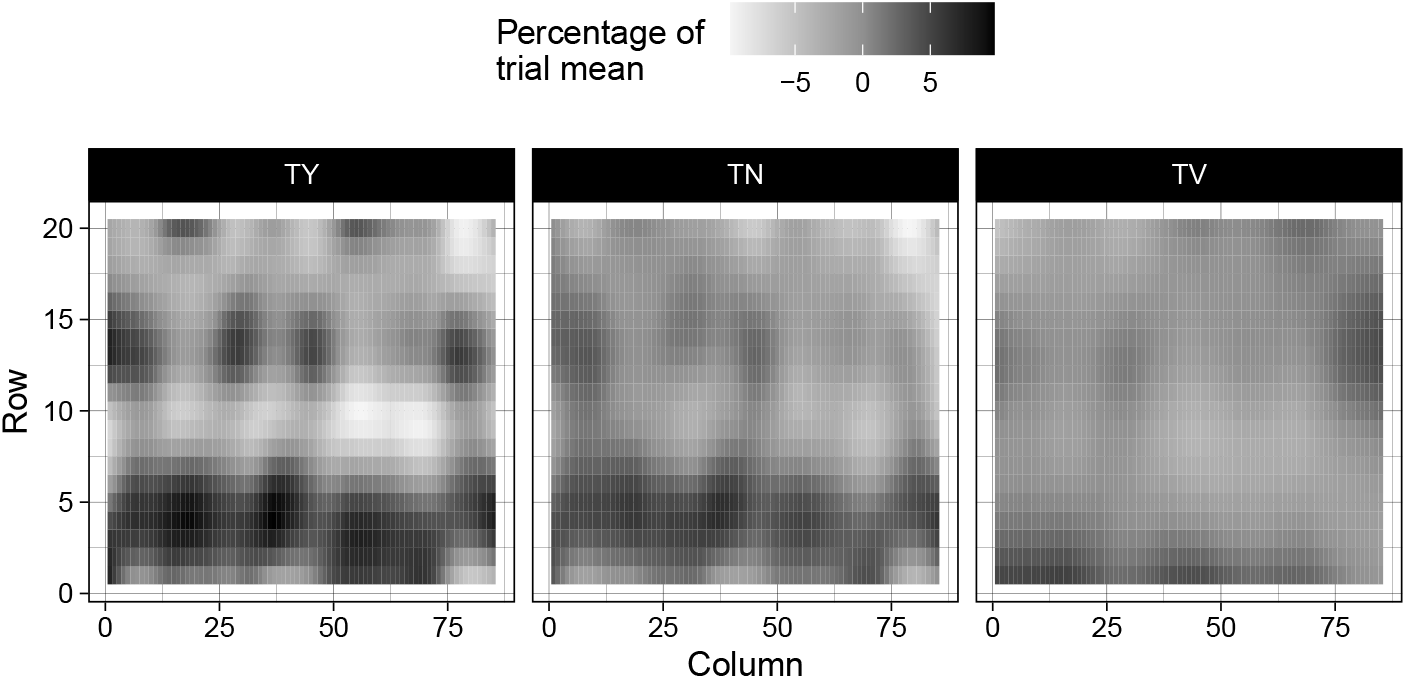
Estimated spatial trends scaled by their trial mean for total yield (TY; 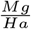), tuber number (TN; Number of tubers per plot), and tuber volume (TV; Average *cm*^3^ per plot) within the Heelsum screening trial

### 2.3 Genetic models

For genetic modelling, F1 hybrids were selected on two criteria: their presence in both screening trials, and whether both parents of a hybrid were utilised in at least two crosses. The former criteria was to ensure estimation of each genotype location combination while the latter was to exclude unconnected crossing sets to guarantee demarcation of parental and cross-wise effects. This resulted in the selection of 225 parental lines which gave rise to 495 F1 hybrid progeny. This panel of hybrids was first utilised in the following multi-location model:

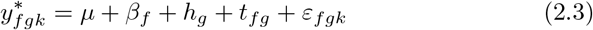

where field location is specified through *β*_*f*_ for the *f*^*th*^ trial location, hybrid performance as *h*_*g*_ for the *g*^*th*^ hybrid, and *t*_*fg*_ is the *g*^*th*^ hybrid by *f*^*th*^ field trial interaction with *ε*_*fgk*_ being an identically and independently distributed residual for the *g*^*th*^ hybrid, *f*^*th*^ field trial, and *k*^*th*^ replicate. From this simple model, best linear unbiased predictions (BLUPs) were made for hybrids within each trial location (*E*[*y*_*fg*_| **h, t**]) as well as over all trials (*E*[*y*_*g*_| **h**]). Each class of conditioned estimates can be found for each trait in Figure 3.

**Fig. 3.**
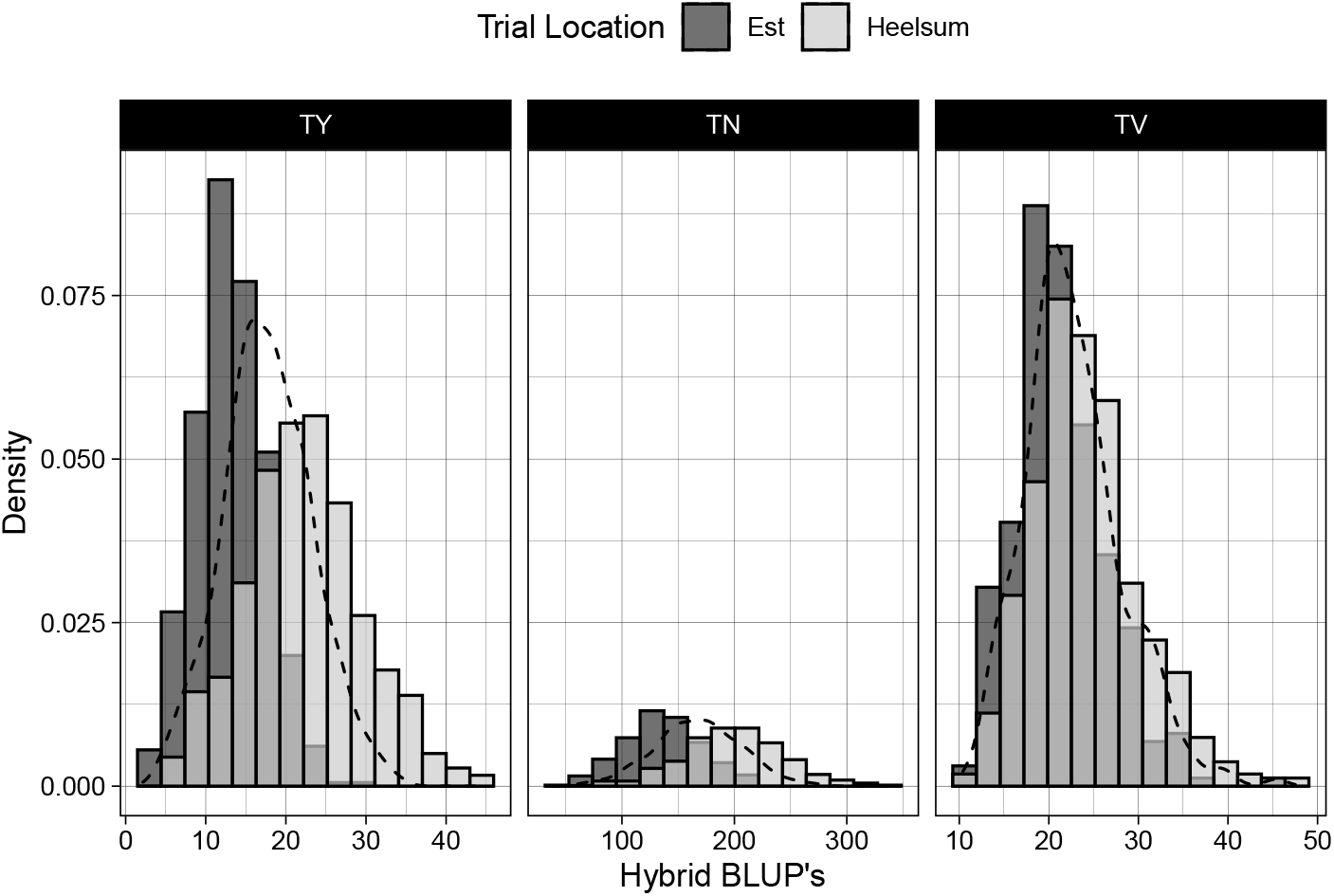
Hybrid best linear unbiased predictions for total yield (TY; 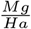) , tuber number (TN; Number of tubers per plot), and tuber volume (TV; Average *cm*^3^ per plot) conditioned on Est and Heelsum trials. Across-location BLUP’s visualised through black density curve.

Of course, the intent of our paper is not merely to retrieve hybrid estimates, but further decompose these estimates into distinct additive and non-additive components. In the context of plant populations, this is traditionally done through a series of controlled crosses between a set of parents which allows for the separation of the parental mean, or the general combining ability (GCA), and the deviation from the expected mean of a cross, or the specific combining ability (SCA) (Sprague and Tatum 1942). These two parameters also have the benefit of being interpreted in terms of genetic variances of a population. The variance attributable to GCA is equal to the covariance between half-siblings while the variance attributed to SCA is equal to the covariance between to full-siblings subtracted by twice covariance of half-siblings (Bernardo 2002). Such models have been made for a variety of population designs (full diallel, half diallel, factorial, etc.) with different effect structures depending on the intent of inquiry. For example, genetic effects can be modelled as fixed if the interest is to provide valid performance estimates for a given cross or they can be treated as random if the variance of effect sampled from a population is desired to be studied (Eisenhart 1947). Additionally, these models can be expanded or simplified accommodating reciprocal effects, trial location or environment interactions, population structures, and so forth. For our purposes, we model in the vein of Griffing’s model II (Griffing 1956) specifying a hybrid’s yield to follow:

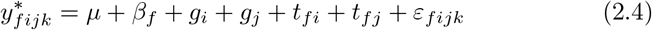

where *μ* being the general mean across trials, *β*_*f*_ is a fixed effect for field trial *f*, *g*_*i*_ and *g*_*j*_ are the GCA’s of parents’ *i* and *j*, respectively, with *t*_*fi*_ and *t*_*fj*_ being their respective field trial and parental interactions with a residual, *ε*, for replicate *k* of the progeny of parents *i* and *j* evaluated in trial *f*. Normal distributions are assumed for all random effects, that is, the maternal and paternal GCA’s 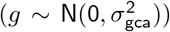, GCA by environment interactions 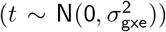, and the resiudal effects 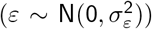 with zero covariance between random effects. This model, containing only the additive main genetic effects, will heron notated as *M*_0_.

This model can be expanded further to include hybrid cross-wise effects with the addition of the SCA term and a SCA by environment interaction. This final model then has the form:

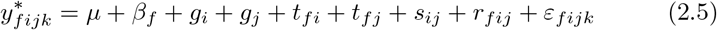

where *s*_*ij*_ is the SCA of parents’ *i* and *j* with *r*_*fij*_ being their respective interaction with trial location *f*. Distributional properties of random effects follow *M*_0_ (Model (2.4)) with the addition of two random effects, those being SCA 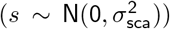 and SCA by environment interaction 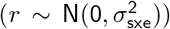 hereon denoted as *M*_*f*_ for all traits going forward. Because the underlying crossing sets are sparse, identifiability of models *M*_0_ and *M*_*f*_ was tested following (Xenakis 2019) to ensure that statistically valid estimates could be derived from all genetic models.

### 2.4 Variance Components

Variance components were estimated for all random effects across all traits studied herein. These variance components were then used to derive several genetic parameters including variation due to additive genetic effects 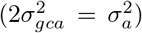, variance due to dominance 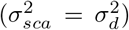 and their respective environmental interactions 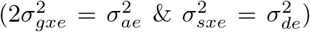. Proportion of total phenotypic variance was then examined with respect to these genetic variances along with each model’s residual variance (Figure 7). Following this, these same genetic components were used to calculate several variance ratios (See Table 2) including broad and narrow-sense heritabilities, 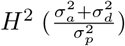 and 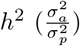, respectively, dominance ratios 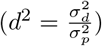, additive portion of genetic variation 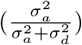, and ratio of additive genetic by environment interaction 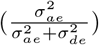. Total phenotypic variation 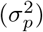 was computed by scaling 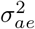 and 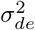 by the total number of field trials and 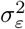 by the product of the number of field trials and the number of replicates used within each trial. Thus, 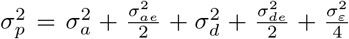. Lastly, realised effects were extracted from the *M*_*f*_ genetic models (equation (2.5)) to derive genetic correlations between traits and trial locations for GCA’s (Figure 4) and SCA’s (Figure 5). Each genetic effect was conditioned on its respective environmental interaction.

**Fig. 4.**
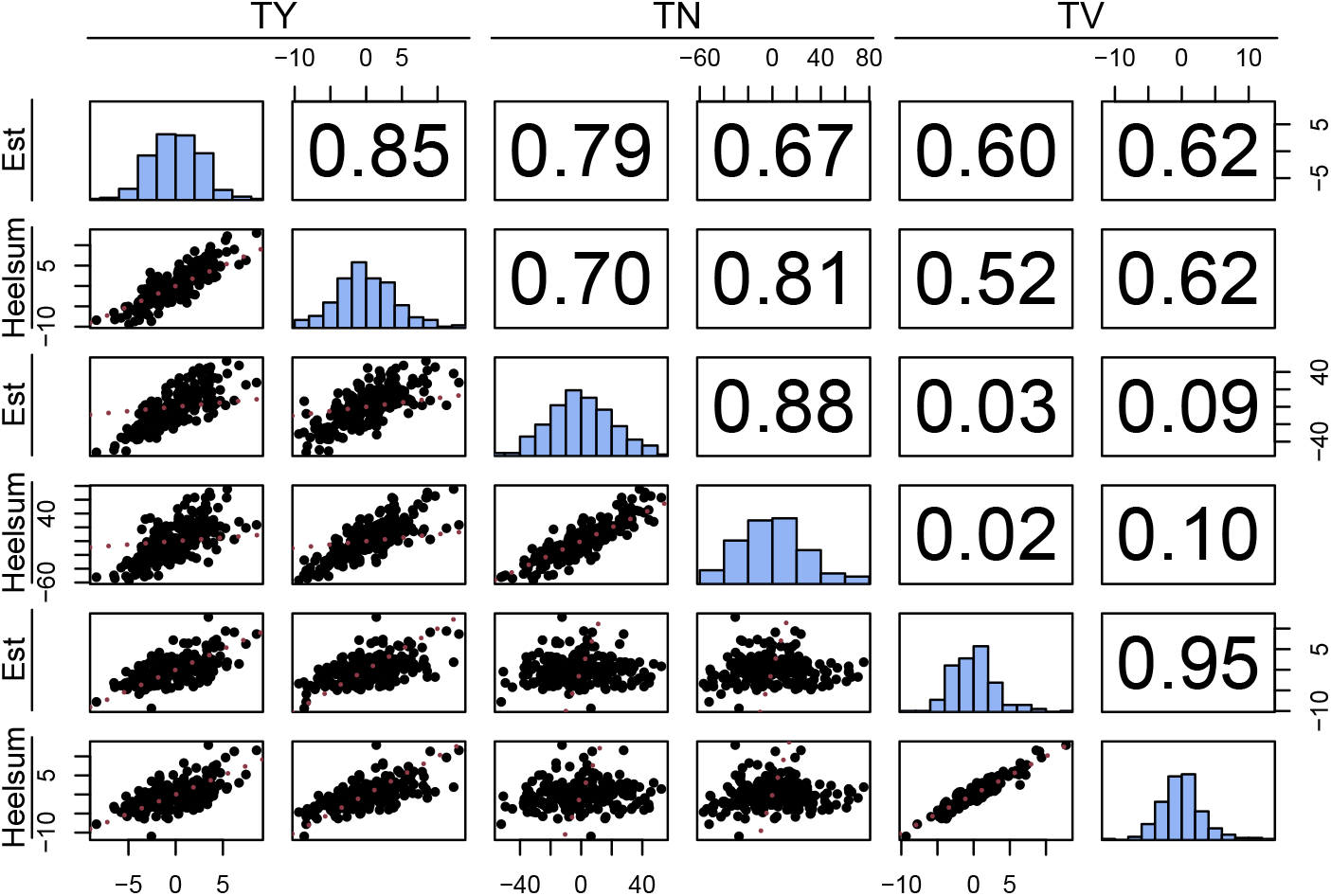
General combining ability pairwise comparisons with scatterplot (lower triangle), Pearson correlation coefficients (upper triangle), and marginal distributions (the diagonal) of best linear unbiased predictions for total yield (TY; 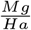), tuber number (TN; total number of tubers per plot), and average tuber volume (TV; *cm*^3^) in Est and Heelsum. The identity is provided in red for each scatterplot (lower triangle).

**Fig. 5.**
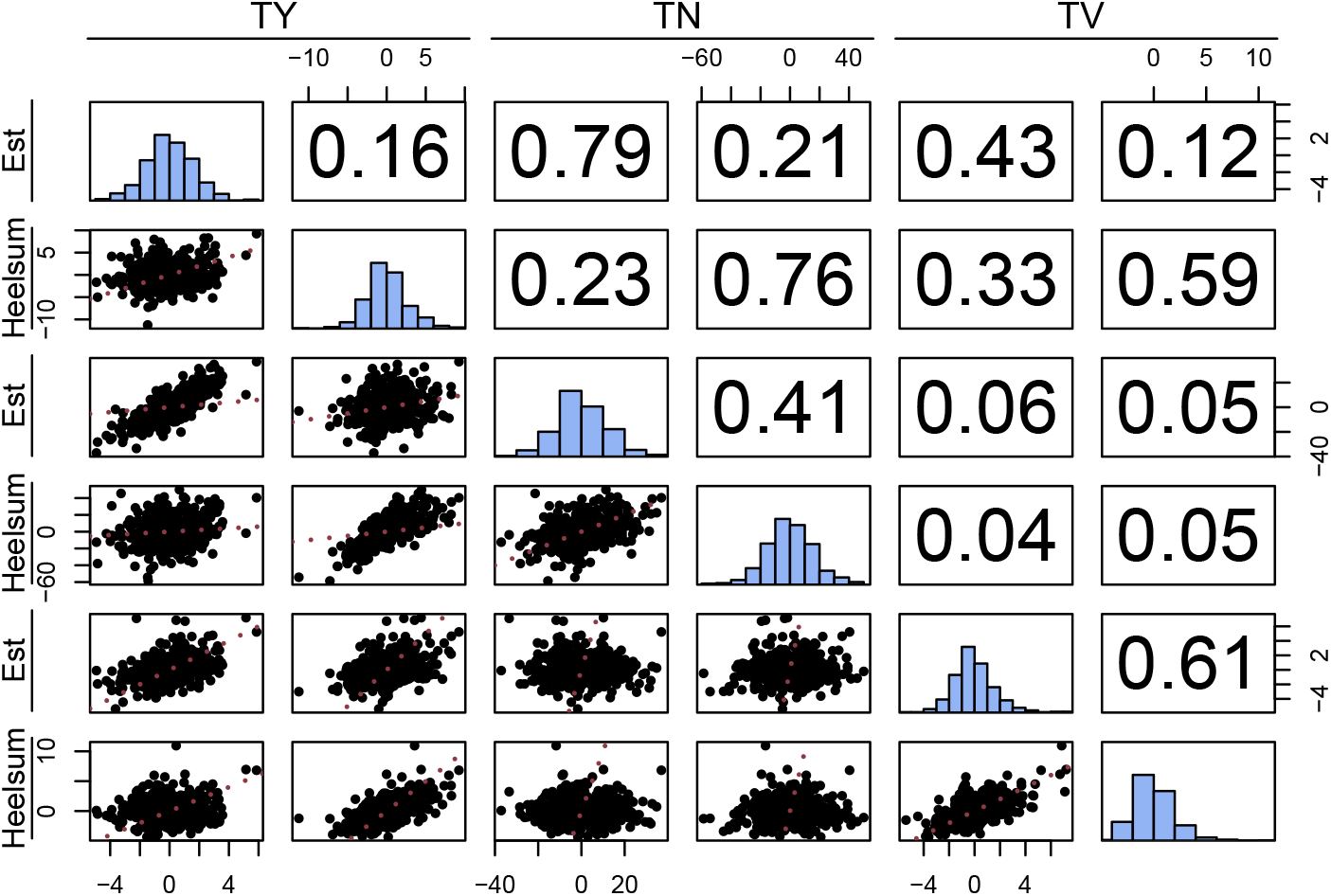
Specific combining ability pairwise comparisons with scatterplot (lower triangle), Pearson correlation coefficients (upper triangle), and marginal distributions (the diagonal) of best linear unbiased predictions for total yield (TY; 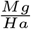), tuber number (TN; total number of tubers per plot), and average tuber volume (TV; *cm*^3^) specific combining ability in Est and Heelsum. The identity is provided in red for each scatterplot (lower triangle).

**Fig. 6.**
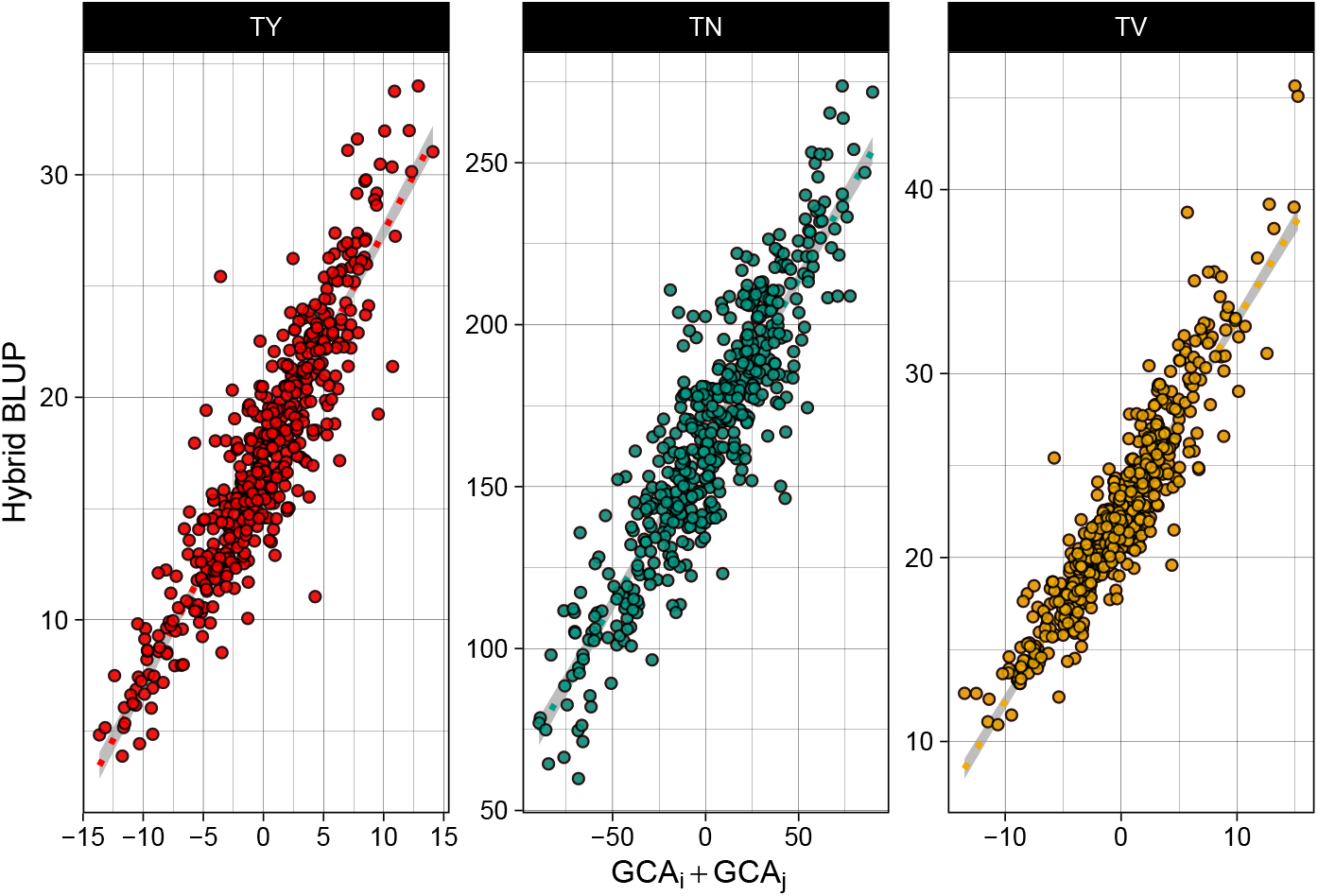
Scatterplot of hybrid BLUP’s regressed on the GCA’s of parents’ i and j for total yield (TY; 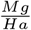), tuber number (TN; total number of tubers per plot), and average tuber volume (TV; *cm*^3^)

**Fig. 7.**
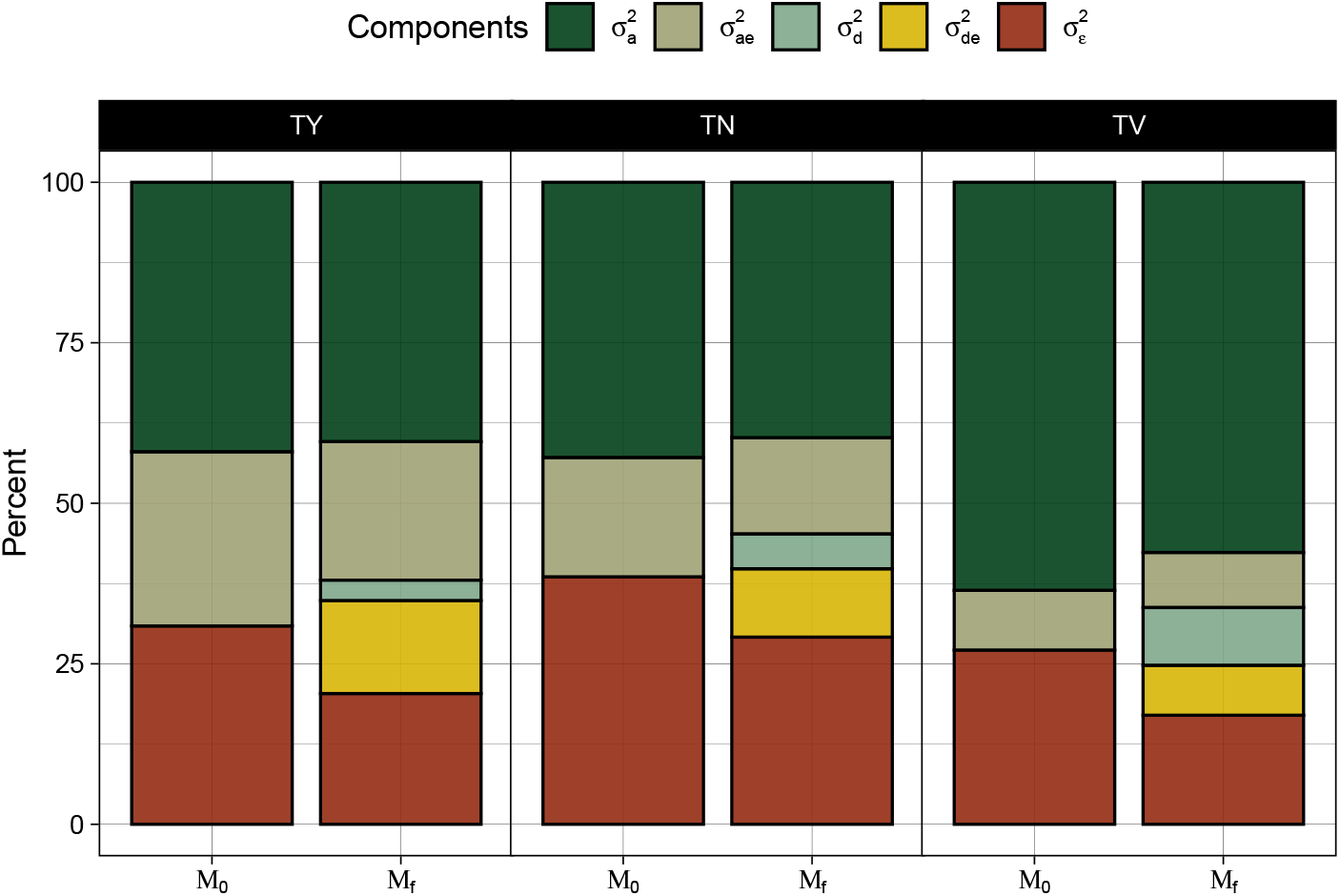
Partitioning of total phenotypic variance into additive 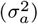, dominant 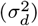, interactive 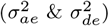, and residual 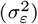 genetic components across each genetic model (*M*_0_ and *M*_*f*_) for total yield (TY), tuber number (TN), and average tuber volume (TV)

**Table 2.** The broad and narrow-sense heritabilities (*H*^2^ and *h*^2^, respectively), dominance ratio (*d*^2^), proportion of additive genetic variation 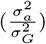, and proportion of additive genotype by environment interaction 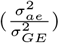 estimated from each *M*_*f*_ models for total yield (TY), tuber number (TN), and average tuber volume (TV)

|  | TY | TN | TV |
| --- | --- | --- | --- |
| $H^2$ | 0.65 | 0.69 | 0.84 |
| $h^2$ | 0.61 | 0.61 | 0.73 |
| $d^2$ | 0.05 | 0.08 | 0.11 |
| $\frac{\sigma_a^2}{\sigma_G^2}$ | 0.93 | 0.88 | 0.87 |
| $\frac{\sigma_{ae}^2}{\sigma_{GE}^2}$ | 0.60 | 0.59 | 0.52 |

### 2.5 Hypothesis testing

To evaluate statistical evidence of heterosis (through SCA term), we perform a simple hypothesis testing procedure where *M*_0_ represents a null model where 270 the additional effects from *M*_*f*_ (i.e. 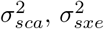) are constrained to zero. Therefore, we can construct a non-standard hypothesis test where:

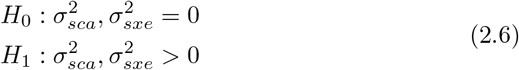

which can be evaluated directly through the following likelihood ratio test where:

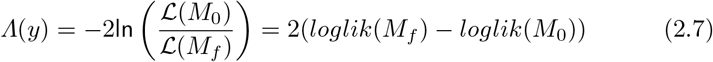

This test is non-standard because it follows a special case where testing is occurring on the boundary of the parameter space which is often taken into account using a mixed *χ*^2^ distribution following (Self and Liang 1987). For our testing purposes, we used a 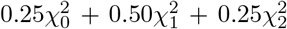 mixture distribution. These tests are also accompanied with the Akaike information criterion (AIC) for each genetic model (Table 3). Under *H*_1_, the hybrid genetic effect from our original model (2.3) should equal the sum of the GCA & SCA effects specified in model (2.5). All models were solved through restricted (residual) maximum likelihood (REML) using ASReml-R 4 (Butler et al. 2017). REML based procedures have come into popular usage over the past two decades due to their ability to provide estimators both consistent and asymptotically normal even under conditions with non-orthogonal sets of random predictors which is particularly useful while using sparse crossing designs (Searle, Casella, and McCulloch 1992). This is not to mention the volume of diallel-based literature where REML is the invoked method of choice for reasons which will not be discussed here ((Möhring, Melchinger, and Piepho 2011) provide an excellent review on the topic). All trial and pedigree data utilised for the following analysis have been made available on GSA FigShare. File `Phenotypes.csv` contains the three aforementioned traits along with field trial row, column, and block indices for each observation. File `Pedigrees.csv` gives a hybrid identification number with each parental code.

**Table 3.** Likelihood ratio tests for *M*_0_ and *M*_*f*_ genetic models together with each model’s Akaike’s information criterion for total yield (TY), tuber number (TN), and average tuber volume (TV)

| | AIC | $\Lambda(y)$ | $P(\Lambda(y))^\dagger$ |
| --- | --- | --- | --- |
| **TY** |  |  |  |
| $M_0$ | 8121 | 165 | *** |
| $M_f$ | 7960 | | |
| **TN** |  |  |  |
| $M_0$ | 15833 | 98 | *** |
| $M_f$ | 15738 | | |
| **TV** |  |  |  |
| $M_0$ | 7089 | 220 | *** |
| $M_f$ | 6873 | | |

$^\dagger$ \*\*\*: $P(\Lambda(y)) < 0.001$

## 3 Results

### 3.1 Spatial Components

Spatial models for the Est and Heelsum trials were created for each yield component. All spatial models successfully converged with credible spatial trends for both trial locations. Model residuals for all environmental models showed little to no evidence of deviation from normality; all of the above suggest successful delineation of spatial components for all traits in both field trials.

Total yield and tuber number exhibited evidence for strong local trends with a row effect contributing to the spatial trend in Est (Figure 1). While these same components also impacted tuber volume, it was not nearly so prominent (See S1). The similar magnitude of row effects on tuber number and total yield can also be observed through the ratio of effective and nominal dimensions which were identical for these two traits (0.68) in contrast to tuber volume (0.47). Additionally, the effective nominal dimension ratio for hybrids (i.e. a generalised heritability) was highest in tuber volume followed by total yield and lastly tuber number (See Table S1).

Overall, the spatial trends in the Heelsum trial were less severe than those evaluated in the screening trial in Est. Interquartile ranges of field effects were between −8.96% and 9.04% of the trial mean for total yield in Est while the range was −2.85% and 3.55% of the trial mean for yield in Heelsum (Figure 2). The differences in magnitude of spatial components between these two trials were also similar for tuber number and tuber volume. Nonetheless, the estimated spatial components did have a modest effect on all yield related components in Heelsum. In particular, random effects for the field column had a minor impact on tuber volume (0.55), tuber number (0.48), and total yield (0.45) (See Table S2). General heritability estimates for Heelsum were akin to those observed in Est with values of 0.9, 0.82, and 0.88 for tuber volume, tuber number, and total yield, respectively.

### 3.2 Hybrid estimates

Using spatially corrected phenotypes from model (2.1), we extracted hybrid BLUP’s from model (2.3) whose frequency is plotted in Figure 3 to visualise gross genetic variation displayed across both trials. Generally, hybrid performance was far greater in Heelsum over Est. The trial mean for total yield was 13.1 and 22.4 *Mg* · *Ha*^−1^ for Est and Heelsum, respectively. Likewise, average tuber number was 136 tubers in Est and 199 tubers in Heelsum. The differences in performance between these two trials coincide with previous trials at these same locations (Dijk et al. 2021). Trial means for tuber volume were comparatively more stable between trial locations with means of 21.3*cm*^3^ in Est and 24.2*cm*^3^ in Heelsum. Along with differences in mean hybrid performance, there was also greater dispersion of phenotypes in Heelsum than in Est. This was especially apparent where total yield in Heelsum displayed a standard error twice as large than total yield in Est. This same marked difference could also be seen in tuber number where BLUP’s in Heelsum exhibited a standard error 1.6 times greater than that which was found in Est. Tuber volume in contrast to the other phenotypes exhibited similar BLUP distributions between both trial locations. Examining these sample distributions in light of the effective and nominal dimension ratios (See Table S1 and S2), it can be safely said that tuber volume showed the greatest stability between trials which coincided with its high generalised heritabilities.

### 3.3 Estimates for GCA and SCA

All GCA and SCA effects (with their respective environmental interactions) were predicted from *M*_*f*_ model. Figures 4 & 5 present these as sets of scatterplot matrices with pairwise comparisons of each genetic effect as well as their individual sample distribution along the diagonal. GCA estimates between trial locations had notable Pearson correlation coefficients for all traits consistent with their narrow-sense heritabilities derived from *M*_*f*_ (Table 2). Also noteworthy, within-trial GCA’s with large genetic correlations were found between total yield and each of its yield components, tuber number (0.79 in Est and 0.81 in Heelsum) and tuber volume (0.60 in Est and 0.62 in Heelsum) according to expectation. All genetic correlations were significant with exception of all pairwise correlations between tuber number and tuber volume (Figure 4). Modest genetic correlations between traits from disparate trials could also be found between total yield and tuber number (0.67 and 0.70) as well as between total yield and tuber volume (0.62 and 0.52).

Similar multivariate trends were observed for realised SCA’s, though, with globally smaller genetic correlations. Within-trait between-trial genetic correlations were modest for the individual yield components (0.61 and 0.41 for tuber volume and tuber number, respectively) while total yield had little correspondence across locations indicative of strong SCA by environment interaction (Figure 5). The pairwise correlations between total yield and the individual components were almost as strong as the GCA correlations; total yield and tuber number had genetic correlations of 0.79 and 0.76 while total yield and tuber volume were 0.43 and 0.59 for Est and Heelsum, respectively. Genetic correlations between tuber volume and tuber number, similar to their GCA counterparts, exhibited no statistical dependence with non-significant correlations in the range of 0.04 and 0.06 (Figure 5).

When comparing the GCA and SCA quantiles for each trait and trial location, the realised GCA effects were consistently larger than those SCA’s estimated. On average, any given GCA quantile was two times larger than its respective SCA quantile; this was true for all traits measured here with some minor departures (See Table S3). These differences in magnitude between the estimated GCA and SCA effects could also be readily seen when regressing hybrid averages on the mid-parent value and full genetic effects (Figure 6). No linear relationship could be found between the estimated GCA and SCA effects for any trait (See Figure S1) coinciding with our model assumptions (2.5).

### 3.4 Variance Components

Variance estimates were derived for all specified random effects for models *M*_0_ and *M*_*f*_. Tuber volume not only exhibited the largest proportion of variance explained by SCA (*d*^2^ = 0.11), but also had the largest total genetic variance of any trait (*H*^2^ = 0.84) (Figure 2). Tuber number harboured some amount of SCA (*d*^2^ = 0.08) but a considerable portion of non-additive effects appeared to be partitioned in the SCA by environment interaction (Figure 7). This was even more pronounced in total yield where most of the non-additive variance was in the in the dominance by environment interaction and not in the main effect (*d*^2^ = 0.05). Broad-sense heritabilities were quite similar between tuber number (0.69) and total yield (0.65) with the main difference being the aforementioned smaller SCA in total yield. In terms of total genetic variance, the additive genetic component was the largest genetic effect across all traits with the ratio of additive genetic variance being highest in total yield (0.93) followed by tuber number (0.88) and tuber volume (0.87). Interestingly, while tuber volume had the smallest estimate of genotype by environment interaction, it also had the smallest proportion of additive genetic by environment interaction (0.52) suggesting that the dominance by environment interaction was just as important in determining the phenotype as the additive by environment interaction. Contrasting variance components between *M*_0_ and *M*_*f*_ shows that the partitioning of 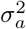 scarcely changes between genetic models suggesting that much of variance from the added effects are being repartitioned from the residual variance (Figure 7).

### 3.5 Model Testing

The likelihood ratio tests conducted between the full genetic model, *M*_*f*_ (containing 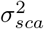 and 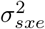, and the null model, *M*_0_, were found to be significant for each trait with each test yielding a probability less than 0.001 (Table 3). Each test conformed with the AIC’s of each model pair where the smallest AIC was observed in the full genetic model suggesting that the best fit was achieved with the inclusion of SCA and its environmental interaction. While *M*_*f*_ gave the best model fit for each trait, there was a marked difference in the value of the test statistic. Tuber volume had the largest test statistic (as well as difference of AIC’s between *M*_*f*_ and *M*_0_) followed by total yield. The test statistic for tuber volume appears to correspond with the size of the SCA variance component in *M*_*f*_ while the inclusion of the SCA by environment interaction appears to have been the predominant component increasing the model fit for total yield.

## 4 Discussion

### 4.1 Presence of additive vs. non-additive effects

SCA was detected across all phenotypes (Table 3) warranting sufficient evidence that SCA can impact yield, especially in its simpler components, seen primarily in tuber volume (Figure 7). However, the magnitude of GCA effects were far greater than the magnitude of SCA’s estimated across all traits irrespective of the their heritability or size of SCA variance. The GCA quantiles were between 1.3 to 2.5 times larger than their respective SCA quantiles (See Table S3) suggesting a systematic importance of the additive genetic effects in this DHP population. The implications then is that most of the variation in the progeny can be found through the additive genetic variation in the parents. This was most apparent for tuber volume (*h*^2^ = 0.73) where regression of hybrid performance on the combination of parental GCA’s had the best fit of all traits (Figure 6). This illustrates that the parental GCA’s (or, identically, the mid-parent value) capture the majority of a hybrid’s phenotype.

While this study is the largest of its kind, it is certainly not alone in attempting to decompose genetic components of yield in potato. A number of similar populations were used in both diploid and tetraploid backgrounds with wide-ranging results. Many of these studies came to find little non-additive genetic variation for yield components similar to the results presented here with GCA being the primary component for traits like average tuber weight, tuber shape, total tuber number, and total yield (Veilleux and Lauer 1981; Brown and Caligari 1989; Neele, Nab, and Louwes 1991). These studies utilised either variants of a diallel or factorial crossing schema making the structure of their statistical models not altogether different than the modelling endeavoured here. One exception between models’ (2.4) and (2.5) and those used in the aforementioned tetraploid populations is that variance attributed to GCA has a different interpretation due to differences in ploidy and levels of inbreeding. This said, there is a large body of work which also finds SCA to be the largest (and at times, the only) detected effect in several complex traits. The same previously mentioned traits as well as others like incidence of hollow heart and tuber uniformity were also found to be predominately under the control of SCA (Killick 1977; Veilleux and Lauer 1981; Thompson and Mendoza 1984; Haynes 2001). Most notably, (G. C. C. Tai 1976) was only able to detect SCA in their partial diallel crosses with no GCA component found for marketable yield and marketable tuber number.

The lack of an empirical consensus on the predominance of GCA and SCA in potato is not very surprising in of itself. The estimation of these parameters are very much contingent on numerous factors including crop ploidy, genetic background, number of parents, degree of environmental stress, and choice of statistical model (to name a few), all of which are prone to change across experiments. Even making comparisons between studies utilising very similar genetic backgrounds can lead to divergent findings (Tarn and Tai 1977; Maris 1989). While seemingly incoherent, the following does offer some interesting grounds for considering those genetic effects observed here. Many of the aforementioned studies estimated variance components on populations which had undergone siginificant selection through a recurrent selection schema (Haynes 2001; Maris 1989) or were themselves the product of strong selection on GCA’s in their ancestors (G. C. C. Tai 1976). In both cases, less additive genetic variation can be expected among hybrids derived from them leaving non-additive genetic variation to be the predominant genetic effect within their inter-population crosses. Conversely, populations like ours which show ample additive genetic variation might be younger with respect to selection pressure in their ancestors on these traits; though without any formal analysis on population structure this is purely speculative. Another line of reasoning for the smaller SCA’s found here relative to many of the tetraploid studies could be explained by *progressive heterosis* whereby higher order non-additive genetic effects become possible through polyploidy (Birchler, Auger, and Riddle 2003). Recent genomic studies in tetraploid potato support this hypothesis with evidence of a genetic residual effect (which could be explained by tri & quadrigenic dominance) contributing as much as 45% of total genetic variation in potato yield (Endelman et al. 2018). However, making any meaningful confirmation on the specific role of ploidy in producing non-additive genetic effects is beyond the scope of this present study and only deserves a cursory mention.

### 4.2 Genetic Architecture of yield

Among our two yield components (i.e. tuber number and and average tuber weight), we found strong genetic correlations between each and total yield for both the additive (see Figure 4) and non-additive genetic effects (see Figure 5). Numerous studies have identified these same phenotypes as major determinants of total tuber yield marking them both as strategic heritable targets for breeding (Khayatnezhad et al. 2011; Thompson and Mendoza 1984). Consistent with these studies, tuber number GCA’s appeared to impact total yield more than tuber volume with *ρ*_TN,TY_ equalling 0.79 and 0.81 and a *ρ*_TV,TY_ of 0.60 and 0.62 in Est and Heelsum, respectively. SCA genetic correlations behaved similarly with *ρ*_TN,TY_ equal to 0.79 and 0.76 and *ρ*_TV,TY_ of 0.43 and 0.59 in Est and Heelsum, respectively. Both the additive and non-additive components point to tuber number being the primary determinant of yield in this hybrid population.

Among certain market classes, our two yield components, average tuber volume (or tuber size) and tuber number, often exhibit an inverted relationship due to the physiological and genetic limits of potato. For example (Thompson and Mendoza 1984) found a genetic correlation of −0.24 among their panel. Additionally, (Lemaga and Caesar 1990) identified negative cubic trends between tuber number and average tuber weight capturing a majority of variation. Interestingly, no meaningful relationship could be found between these two yield components with respect to additive (Figure 4) and non-additive genetic correlations (Figure 5). To repeat our previous suspicion, this suggests a lack of directional selection on one of these two traits evident by the lack of genetic constraints between them (Blows and Walsh 2009). Tuber volume had the largest proportion of additive variation (Figure 7) and genetic variation in general (Table 2) suggesting little to no direct selection on this component of yield. These properties then make this population an interesting candidate for future selection given its genetic potential to be adapted to a variety of tuber types. Another oddity to consider here is that while SCA was detected independently in these two yield components, this did not manifest in the increase of SCA in total yield, but very much the opposite. Others have identified that heterosis in these same yield components were responsible for a geometric increase in gross yield (Tarn and Tai 1977). Further multivariate studies of vigour in diploid potato could further elucidate SCA architecture especially as selection pressure is applied, a key scenario within breeding programmes.

### 4.3 Using GCA & SCA in commercial breeding

The large narrow-sense heritabilities and magnitude of the GCA’s found here have major implications for breeders of DHP. To begin, the valid estimation of GCA’s further validate the potential of potato in its conversion into a inbred-hybrid crop as purported before (Jansky et al. 2016; Lindhout et al. 2011). Further, the size of the estimated GCA’s relative to a hybrid’s average performance (Figure 6) show that standard breeding designs used in other hybrid crops will likely be just as efficacious in DHP. For example, the use of test crosses, a mainstay in maize breeding, can also be utilised in evaluating performance of potato parental lines for hybrid crosses with reasonable success. These test crosses can be further utilised for model training and be the basis for genomic selection of parents (using breeding values) or hybrid crosses (via mid-parent value); again, a standard-place method in hybrid breeding (Albrecht et al. 2011). To quickly add, while we found little contributions made by SCA, their relevance to breeders do not necessarily end here. Depending on the specific mechanism behind these observed non-additive effects, they could be further exploited in the trait of interest through initial breeding design (e.g. formation of heterotic pools). Heterotic breeding has become a major target for quality trait improvement in other solanaceous crops including chilli pepper (Herath et al. 2020), eggplant (Kumar et al. 2020), and tomato (Frankel 1983). This is where potato meets an interesting intersection between the vegetable and agronomic worlds where SCA might play a more valuable role for qualities controlling specific market criteria (e.g. average tuber volume, tuber length & shape) but would be less emphasised in composite and complex traits (e.g. gross yield, starch & protein content) where GCA is the predominant genetic effect at play. Having said this, future work into the biometric mechanism of vigour will be able to lend more wisdom for how breeders should wield this in a hybrid potato breeding programme.

### 4.4 Future work

Finding evidence for heterotic effects in DHP does not yield much regarding the source of the effects identified here. The statistical models assume all underlying effects captured by the SCA term to be the product of cumulative dominance deviations across the genome, but there exists many other plausible sources of non-additive variation. Since its conception, genetic theory has explained heterosis with a whole suite of models with many being broadly plausible (See (Labroo, Studer, and Rutkoski 2021)). However, these apparent non-additive effects could just as simply be explained by dispersion of additive alleles among parents, a hypothesis which is generally supproted emperically (Frankel 1983; Mackay et al. 2021). Nevertheless, these effects continue to be interesting point of study and still remain an important target in hybrid breeding of modern crops. This is especially worthwhile in DHP given its novelty as a hybrid crop with the potential of heterotic breeding to still be determined.

## 5 Conclusion

This is the first study in diploid hybrid potato to produce estimates of general and specific combining abilities using a large panel of commercially-derived paorents. This represents a major milestone in the reorientation of potato from a clonal tetraploid to a diploid inbred-hybrid crop. Identifying the predominance of additive genetic effects for multiple yield components among hybrids offers strategic insight on the necessity of effective generation of parental lines and early population development in general. Though the estimated non-additive effects in this population are smaller in contrast to their additive counterparts, heterotic vigour shows some minor role in simpler traits. Specific quantitative traits should be targeted for SCA exploitation to bolster variety development on top of their parental effects. Further research into the genetic mechanisms for the apparent non-additive effects will also better elucidate the strategic advantage (if any exist) in key economic targets in hybrid potato.

## 6 Statements and Declarations

### 6.1 Competing Interests

On behalf of all authors, the corresponding author states that there is no conflict of interest.

### 6.2 Author Contributions

**JR Adams**

The primary author and researcher in the composition of this manuscript.

**ME Vries**

Domain expert and researcher who offered relevant guidance regarding the larger relevance to potato.

**C Zheng**

Supervisor and researcher who helped in the earlier stages of statistical modelling and assisted in the communication of the methods used here.

**FV Eeuwijk**

Primary supervisor who helped verify the methods used in this manuscript and confirmed the validity of the results presented here and their implications.

### 6.3 Key Message

Hybrid vigour was detected for multiple traits in diploid hybrid potato. Additive gene action was most prominent in tuber yield and should be the primary target within hybrid breeding programmes.

